# Efficacy and accuracy responses of DNA mini-barcodes in species identification under a supervised machine learning approach

**DOI:** 10.1101/2020.12.10.420281

**Authors:** Mohimenul Karim, Rashid Abid

## Abstract

Specific gene regions in DNA, such as cytochrome c oxidase I (COI) in animals, are defined as DNA barcodes and can be used as identifiers to distinguish species. The standard length of a DNA barcode is approximately 650 base pairs (bp). However, because of the challenges associated with sequencing technologies and the unavailability of high-quality genomic DNA, it is not always possible to obtain the full-length barcode sequence of an organism. Recent studies suggest that mini-barcodes, which are shorter (100-300 bp) barcode sequences, can contribute significantly to species identification. Among various methods proposed for the identification task, supervised machine learning methods are effective. However, any prior work indicating the efficacy of mini-barcodes in species identification under a machine learning approach is elusive to find. In this study, we analyzed the effect of different barcode lengths on species identification using supervised machine learning and proposed a general approximation of the required length of the minibarcode. Since Naïve Bayes is seen to generally outperform other supervised methods in species identification in other studies, we implemented this classifier and showed the effectiveness of the mini-barcode by demonstrating the accuracy responses obtained after varying the length of the DNA barcode sequences.

## I. Introduction

Species identification is an elementary component in biological research. According to the Catalogue of Life, a hierarchically ranked system based on expert opinions provided by taxonomists, more than 1.6 million species have been identified to date [1]. However the total number of species is estimated to be 8.7 million [2]. Therefore, despite over 250 years of taxonomic classification, the existing records cover only a small portion of species on Earth (14%) and in the ocean (9%) with the remaining yet to be described [2]. From a taxonomic perspective, species discovery comprises three pivotal and onerous steps: specimen collection, species level sorting, species identification/description [3]. These steps consider morphological characteristics in the traditional ‘evidence-based’ taxonomic approach and cause challenges in the decision-making process [4]. Morphological characteristics can be significantly different for the same species in different stages of its life-cycle. In case of archival museum specimens, distinguishing morphological details are scarce. During the specimen collection phase organism fragments, stomach contents, feces, saliva, skin-scraps, blood, and pollens are collected, which often lack substantial morphological characteristics. Some congeneric, species-complex and cryptic species have only tiny morphological differences, and thus are often misclassified as a single species. In sexual dimorphism, different sexes of the same species can exhibit significantly disparate morphological attributes, leading to misidentification as different species.

To address all the aforementioned issues with classification using traditional morphological methods, a more advanced, accurate, fast and efficient alternative approach is required to attenuate morphological dependencies. A microgenomic approach involving DNA sequences has been established recently as a promising technique for species identification [5]. In this technique, a short portion of DNA from a specific gene of an organism is used to uniquely identify a species in comparison to a reference library [6]. This specific portion is called the DNA barcode of that species. Different gene regions are used as DNA barcodes for different groups, such as the cytochrome-c-oxidase-I (*COI/COX1*) gene found in the mtDNA of animals [7], *rbcL* and *matK* in plants [8], and the internal transcribed spacer (*ITS*) rRNA in fungi [9]. Species identification using DNA barcodes can be considered as a classification problem where an unknown DNA barcode sequence is matched against a reference library of already known species DNA barcode sequences [10]. Several methods have been proposed to assist with this classification matching [11] with the supervised machine learning approach being prominent. In this approach, the reference dataset of known species is used as the train set for training the corresponding machine learning models and the barcode sequences requiring classifications are placed in the test set for the query [12].

Notwithstanding the prominence of DNA barcode in species identification, several shortcomings arise. Although polymerase chain reaction amplification and sequencing are usually consistent in freshly collected and well-preserved specimens, obtaining a full length (~650 bp) barcode in archival museum specimens that are preserved under sub-optimal or DNA-unfriendly conditions is difficult [13]. This is also the case for processed biological materials and decayed tissues owing to the degraded quality of the DNA [14]. In cases where obtaining a full-length barcode is difficult, shorter barcode sequences (~ 100–300 bp), often referred to as *mini-barcodes*, are useful for species-level identification owing to their comparatively better amplification gain [15]. Mini-barcodes are also commonly used in DNA metabarcoding strategies based on environmental DNA [16] and for studying ecological biodiversity [17]. Mini-barcodes can also be applied in alternative next generation short-read sequencing platforms providing higher throughput while remaining cost-effective [13].

## II. Related Works

DNA Barcodes were proposed as an effective molecular approach in species diagnosis by Hebert *et al.* [6] and have since been used in a wide range of studies [18]–[20]. The International Barcode of Life and the Consortium for the Barcode of Life have supported development initiatives for establishing DNA barcodes as a global standard for species identification [21]. Species identification approaches using DNA barcoding can be partitioned into a few major categories: distance based approach, similarity based approach, phylogenetic approach, statistical approach, and character based approach [11], [22], [23].

In distance based approaches, the reference library sequences are ranked with respect to their distances with the query barcode sequence, as small distances suggest that two barcodes can be of specimens of the same species whereas large distances indicate that they belong to two different species. This distance, commonly referred to as barcode gap [24], can be calculated based on the pairwise p-distances [6] or K2P distance [22]. TaxI [25] uses pairwise distances as distance measures whereas BOLD [26] uses K2P distances by default. In similarity based approaches, the query sequences are assigned to species based on a similarity score [27] indicating similarities in barcode characters. BLAST [28], FASTA [29], TaxonDNA [30], NN [22] are some prominent examples in this regard. Phylogenetic approaches construct trees using hierarchical clustering [27] where distinct clusters represent distinct species [31]. Phylogenetic approaches can adopt NJ [32], parsimony (PAR [33]), maximum likelihood (PhyML [34]) or bayesian inference (MRBAYES [35]) as underlying methods. Statistical approaches use population genetics assumptions [36] based on coalescent theory (GMYC [37]) considering phylogenetic uncertainty (SAP-NJ [38]), likelihood ratio tests [39] etc. Character based approaches investigate the presence/absence of particular characters, known as diagnostic sites, instead of total sequence imitating morphological approaches [23]. CAOS [40], BLOG [10], DNABar [41], BRONX [27], DOME-ID [27] are some instances of this approach. There are alignment-free tools such as: ATIM-TNT (Tree-based) [27], CVTreeAlpha1.0 (Component Vector based) [42], Spectrum Kernel [43] and Alfie (python based) [44]. Web Based Tools such as: Linker [45], iBarcode [46], BioBarcode [47] and ConFind [48] are also there for DNA barcoding. Extensive comparative analyses of these major classical barcoding approaches have been studied [11], [27]; however, there is no general consensus on a single optimal method to analyze DNA barcode data for species identification [22], [23].

Machine learning methods have been introduced as state-of-the-art techniques for their extensive applicability in species identification using DNA barcoding [12]. Although supervised statistical classification methods such as: Classification And Regression Tree, Random Forest, and Kernel methods have been studied previously [22], supervised machine learning algorithms e.g., SVM, Jrip, J48 (C4.5), and Naïve Bayes were compared on WEKA platforms to the ad-hoc methods by Weitschek *et al.* and SVM and Naïve Bayes outperformed the other methods for synthetic and empirical datasets [12]. Simple-logistic, IBK, PART, attribute-selected classifier, and bagging approaches were also implemented in [49]. SMO, BP-NN [50], RF [51], and k-mer based approaches [52], [53] can be seen in recent studies. Naïve Bayes was applied on the COI [54] barcode database to show misclassification rates and on ribosomal databases [55].

With the exploration of the prospects of DNA minibarcode in species identification [56], the application of DNA barcode significantly broadened. It is observed that although full length barcodes perform efficiently with 97 % species resolution, 90 % and 95 % identification success can be obtained with 100 bp and 250 bp mini-barcodes, respectively [15]. In case of species diagnosis from damaged/degraded DNA samples, ~ 100–300 bp mini-barcodes have shown promising results [14]. Tree-based methods, objective clustering, automatic barcode gap discovery, and Poisson tree process were examined to analyze the use of mini-barcodes for species identification [3]. No simple formula exists to determine the necessary sequence lengths for species identification [7] but the expectation of achieving significantly better performance using full-length barcodes is questionable because a saturation point can be obtained beyond which the additional length data yield insignificant impact compared to the increased overhead costs [3]. In the present study, we examined this issue from the machine learning perspective by demonstrating the accuracy responses of species identification at different partial barcode lengths after using a Naïve Bayes classifier.

## III. Datasets

### A. Empirical data

For species identification, a full reference set with all possible nucleotide polymorphisms for the sequences of each species is required and an adequate reference set is important to prevent over-fitting and under-fitting [12]. Therefore, we included the datasets used in [12] in our analysis, which are a collection of published empirical datasets and simulated DNA barcode datasets. The public empirical datasets of *Bats, Birds, Cypraeidae, Drosophila*, and *Fishes* were selected from the GenBank Nucleotide Database for high phylogenetic diversity and are available for download at^1^ in FASTA format [12]. The datasets referred to here as *Bird2, Fish2*, and *Butterfly2* were extracted from BOLD and are available in CSV format at ^2^. An additional dataset considered in our analysis is the first large-scale genetic assessment of the butterflies of Southern South America (available for download at ^3^) showing several cases of both deep intraspecific and shallow interspecific divergence [57]. The *COI* gene was used as the gene region in the DNA barcode sequences for all these datasets. A detailed overview of these empirical datasets is presented in Table I.

**TABLE I.** Summary of the Empirical Datasets: COI Gene as Barcodes.

| Dataset | No of sequences | Sequence Length | No of Species | Gene Region | Ref |
| --- | --- | --- | --- | --- | --- |
| Cypraeidae | 2,008 | 614 | 211 | COI | [58] |
| Drosophila | 615 | 663 | 19 | COI | [59] |
| Bats | 826 | 659 | 82 | COI | [26] |
| Fishes | 626 | 702 | 82 | COI | [60] |
| Birds | 1,700 | 691 | 150 | COI | [61] |
| Bird2 | 2589 | 990 | 656 | COI | [62] |
| Fish2 | 754 | 901 | 211 | COI | [63] |
| Butterfly | 2028 | 658 | 417 | COI | [57] |
| Butterfly2 | 4267 | 901 | 561 | COI | [56] |

To investigate the overall scenario of species identification using DNA barcoding and analyze the accuracy responses with varying barcode lengths, we used datasets of plants and fungi as well. These empirical datasets use the *rbcL* and *ITS* gene regions, respectively and are available for download at ^1^ in FASTA format [12]. A detailed overview of these empirical datasets is presented in Table II.

**TABLE II.** Summary of the Empirical Datasets of Fungi, Inga, and Algae.

| Dataset | No of sequences | Sequence Length | No of Species | Gene Region | Ref |
| --- | --- | --- | --- | --- | --- |
| Fungi | 50 | 930 | 8 | ITS | [9] |
| Inga | 913 | 1,838 | 56 | ITS | [64] |
| Algae | 26 | 1,128 | 5 | rbcL | [8] |

### B. Simulated data

We used simulated DNA barcode datasets for additional analysis. These datasets are available for downlaod at ^1^ in FASTA format [12]. As stated in [11], these real DNA barcode datasets were simulated considering the time of species divergence and the effective population size (*N_e_*). To simulate gene trees *N_e_* = 1000, 10000, and 50000 was used and the dataset complexity was increased with population size. Thereafter, DNA barcode sequences were simulated on the additive gene trees with 650 base sequence-length with similarity to the real-life size of a standard DNA barcode. Each dataset consisted of 50 species and 20 individuals per species. A detailed overview of the simulated datasets is presented in Table III.

**TABLE III.** Summary of the Simulated Datasets.

| Dataset | $N_e$ | No of Individual | Sequence Length | No of Species | Ref |
| --- | --- | --- | --- | --- | --- |
| Ne1000 | 1000 | 20 | 650 | 50 | [11] |
| Ne10000 | 10000 | 20 | 650 | 50 | [11] |
| Ne50000 | 50000 | 20 | 650 | 50 | [11] |

## IV. Methodology

### A. Naïve Bayes

Naïve Bayes [65] is a method that is frequently used if a large reference set is available. Naïve Bayes is based on Bayes’ theorem, and it is assumed that the predictors are independent of each other. If an instance S with n features is represented by a feature vector *x* = {*x*_1_, *x*_2_,…, *x_n_*}, the model assigns a probability *P*(*C_m_*|*x*_1_, *x*_2_,…, *x_n_*) to S. Here, *C_m_* is the *m^th^* class among all possible classes.

Using Bayes’ theorem, we calculated the posterior probability *P*(*C_m_*|*x*) from the likelihood *P*(*x*|*C_m_*), prior probability *P*(*C_m_*), and evidence *P*(*x*):

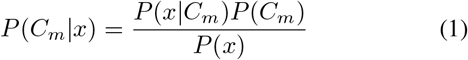

Under naïve conditional assumption,

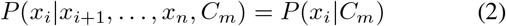

Therefore, the posterior probability *P*(*C_m_*|*x*) was calculated as follows:

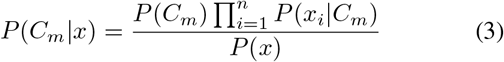

Because *P*(*x*) is constant for a given input, we can only consider the numerator.

Using the Naïve Bayes model, the classification rule can be implemented as follows, where 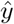 is the class label.

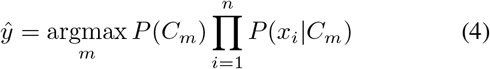

### B. Species identification using Naïve Bayes

Because of the efficacy of Naïve Bayes algorithm in supervised prediction problems [54] and its better performance compared to other supervised machine learning methods in species identification [12], we have used this model for species identification process. The identification approach used was as follows:

Let *b_k_* be the *k^th^* barcode sequence of length *n* in the dataset where *k* = 1, 2,…, *n_b_* and *n_b_*= no of barcode entries in the dataset. *x*_1*k*_, *x*_2*k*_,…, *x_nk_* are the nucleotides in the *b_k_* barcode sequence. If *C_p_* is the *p^th^* species among the different classes of species, the probability that the barcode sequence *b_k_* belongs to the species *C_p_* is

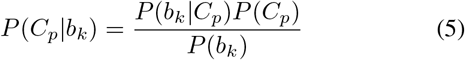

The prior probability in eq. 5 can be calculated as follows:

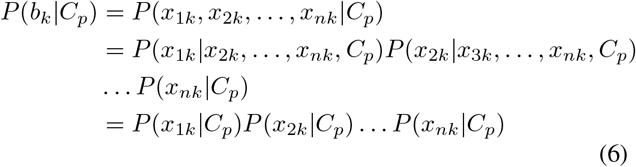

In the above equation, *x_ik_* refers to the nucleotide base at the *i^th^* position of *k^th^* sequence in the dataset and *P*(*x_ik_*|*C_p_*) means the probability of occurring nucleotide base *x_ik_* in species *C_p_* at position *i* where *i* = 1,…, *n*. Therefore, if *N_C_p__* and *N_x_ik__* are the number of samples of *C_p_* and the number of occurrences of *x_ik_* at position *i*, respectively, then we calculate this probability as

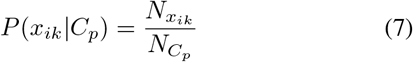

If the proportion is equal to 0, then the posterior probability becomes 0. To prevent this, we replaced zero with a significantly small fractional value *α*, where 0 < *α* << 1. This *α* can be considered as the mutation rate of nucleotides in the particular gene region [54]. *α* = 9.7 × 10^−8^ was proposed as the estimated value of this mutation rate in COI genes [66] and hence it is also used here. For ambiguous bases (e.g., *N*) or missing data (e.g., -) at any position, we have not considered those positions for identification purposes. In some datasets, there is a significant number of missing data (i.e., -) in a significant number of sequences because of sequence alignment. Ignoring these missing data in such cases could affect the performance of our model during the identification process. Therefore, in these situations, we replaced a missing value in a particular position with the most frequently occurring nucleotide in that position within a species.

### C. Data preparation

The empirical and simulated DNA barcode datasets discussed in the section III were used to perform our analysis. Most of these empirical datasets were partitioned by biologists into 80 % and 20 % of total sequences per species in training and testing sets, respectively, maintaining the necessary polymorphism and species divergences [11]. In cases where only a single dataset file was provided, it was partitioned into train and test sets based on an 80 %-20 % random split. However, each dataset consisted of four or more representing sequences per species in the train set where possible, since it was necessary for obtaining a reliable classification performance [12]. The simulated datasets were also split into train and test sets in a 80 %-20 % ratio where for every species 16 sequences were placed into the train set and 4 sequences were placed into the test set. In cases where the datasets were non-aligned, MAFFT [67] was used as a multiple sequence alignemnt tool for aligning the barcode sequences. Most of the datasets were in FASTA format and a Python program was written to convert these FASTA files into corresponding CSV files. For large numbers of missing nucleotides in the barcode sequences of a species, the most frequently occurring nucleotide within that species was considered as the replacement for that missing value. The related Python scripts written for converting the files to the desired CSV format and for replacing missing values are available at ^4^.

### D. Experimental arrangement

For species identification with DNA barcodes, we developed a Naïve Bayes classifier using Java (available at^4^). During the training phase, a probability matrix for each known species is calculated. To achieve the goal, a Hashmap was used where the key was the species name and the value was the probability matrix. The matrix stored the probability of each nucleotide base at each position in the barcode sequence for a species. A worked-out example is presented in Table IV where there are four different species and each has three different barcode sequences of length five. The resulting probability matrix of species 1 is presented in Table V.

**TABLE IV.** Sample Barcode Sequences of Four Species.

| Species 1 | Species 2 | Species 3 | Species 4 |
| --- | --- | --- | --- |
| ATTGC | TCCAG | GTATG | CGAAT |
| ATTCC | TCCGG | GCATG | CGGAT |
| ATTGC | TCCGC | GTATC | CGAAT |

**TABLE V.** Probability Matrix of Species 1.

| Bases | Position 1 | Position 2 | Position 3 | Position 4 | Position 5 |
| --- | --- | --- | --- | --- | --- |
| A | 3/3 | 0/3 | 0/3 | 0/3 | 0/3 |
| T | 0/3 | 3/3 | 3/3 | 0/3 | 0/3 |
| G | 0/3 | 0/3 | 0/3 | 2/3 | 0/3 |
| C | 0/3 | 0/3 | 0/3 | 1/3 | 3/3 |

During the test phase, for a query sequence, we calculated the likelihood of this being a particular species. Our model calculated the maximum likelihood and classified the new sequence to that corresponding species. For example, if we obtained a query sequence TCCAC from the test set, our model took the probability of T, C, C, A, C in position 1, 2, 3, 4, and 5, respectively, from the probability matrix of species 1, 2 and so on. If the highest probability value considering the full length of query sequence came from the matrix of species 2, the sequence was classified as species 2. Figure 1 shows an overview of this classification process.

**Fig. 1.**
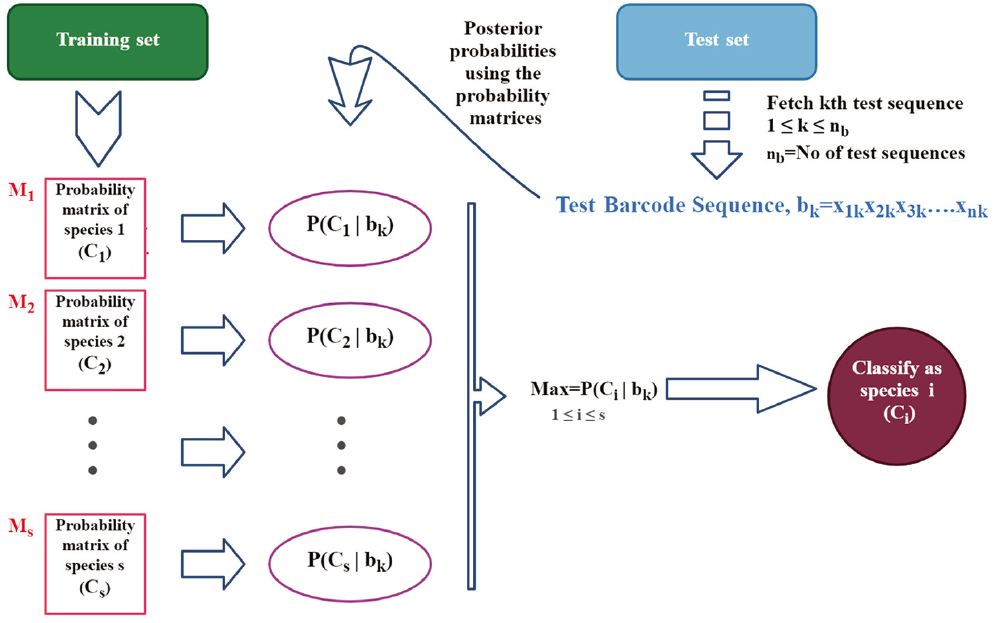
Overview of species identification with Naïve Bayes procedure. Each test sequence is fetched from the test set and using the probability matrices the posterior probabilities are calculated. Based on the maximum probability score, that respective test sequence is classified to corresponding species label.

To analyze the influence of DNA barcode sequence length on species identification, we ran our model with different sequence lengths on all the empirical and simulated datasets.

## V. Results and discussions

After implementing our Naïve Bayes classifier, we tested the efficacy of our method. To ensure the efficacy of our model, we ran our Naïve Bayes classifier on the first five of the COI gene-based empirical datasets presented in Table I considering their full length barcode sequences. By comparing the results to the maximum accuracy from related studies [12], [49], where comparative performance analysis of different supervised machine learning approaches on those same COI gene-based empirical datasets are provided, we found our Naïve Bayes classifier was competent in its performance (as shown in Figure 2).

**Fig. 2.**
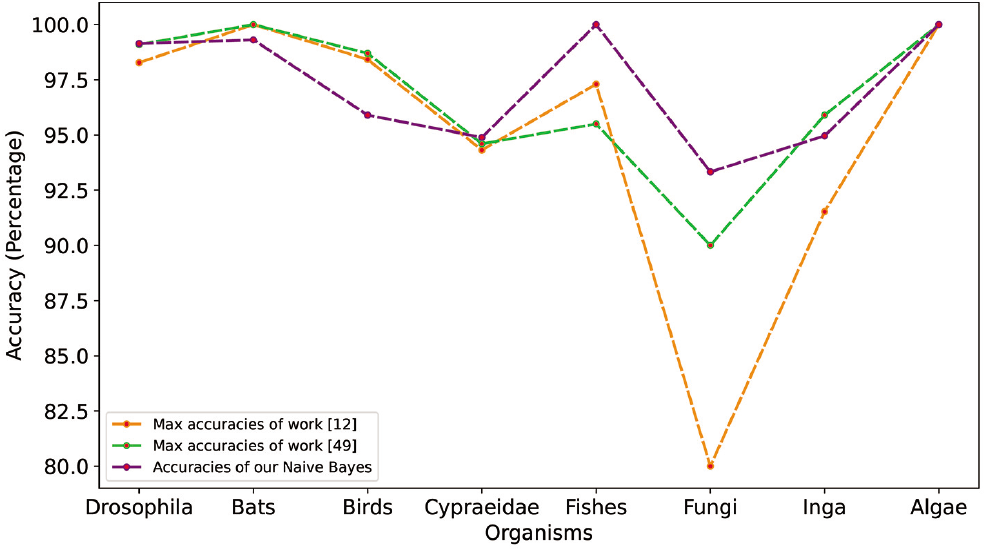
Performance competency of our Naïve Bayes classifier. The accuracy comparison of our model and the maximum accuracy score giving models of the works of [12] and [49] is depicted in this figure. Our model yields very competing performance in each cases.

After testing our Naïve Bayes classifier on the first five empirical COI-based datasets, we ran our model on all of the COI-based empirical datasets presented in Table I to analyze the accuracy responses while varying the barcode sequence lengths. As the datasets in Table I had barcode sequence lengths >600 bp, we set the upper bound of sequence length at 600 while varying the lengths. The sequence lengths were then decreased from the upper bound 600 bp in steps of 50 bases and the corresponding accuracy scores were calculated in each case. As 50 bp was too low a sequence length, we considered a base length of 70 after 100 bp and maintained this as the lower bound for varying sequence lengths in the present study (with a few exceptions). The accuracy scores for the different DNA barcode lengths for the COI-based empirical datasets are presented in Table VI. These lengthwise accuracy scores in Table VI are plotted in Figure 3 and demonstrated in the heatmap in Figure 4. From the pattern of these lengthwise changes of accuracy scores in these plottings and heatmap, it seems that in most cases the accuracy scores remained persistent in the range of 200-600 bp. Although there was a gradual but tiny decrease in accuracy in this range with the decrease of barcode sequence length, it was significantly reduced below 200 bp.

**TABLE VI.** Accuracy Responses in the COI-based Empirical Dataset.

| Datasets | Lengths |  |  |  |  |  |  |  |  |  |  |  |
| --- | --- | --- | --- | --- | --- | --- | --- | --- | --- | --- | --- | --- |
|  | 70 | 100 | 150 | 200 | 250 | 300 | 350 | 400 | 450 | 500 | 550 | 600 |
| Bats | 93.06 | 97.22 | 99.31 | 99.31 | 99.31 | 99.31 | 99.31 | 99.31 | 99.31 | 99.31 | 99.31 | 99.31 |
| Bird | 90.54 | 97.48 | 98.42 | 98.42 | 98.42 | 97.48 | 97.16 | 97.48 | 97.16 | 97.16 | 97.16 | 97.16 |
| Cypraeidae | 89.77 | 91.19 | 93.75 | 94.60 | 94.03 | 94.60 | 95.17 | 95.17 | 95.17 | 95.45 | 95.45 | 95.45 |
| Drosophila | 96.55 | 98.28 | 98.28 | 98.28 | 98.28 | 99.14 | 99.14 | 99.14 | 99.14 | 99.14 | 99.14 | 99.14 |
| Fish | 88.29 | 93.69 | 99.1 | 100.0 | 100.0 | 100.0 | 100.0 | 100.0 | 100.0 | 100.0 | 100.0 | 100.0 |
| Butterfly | 89.8 | 93.47 | 95.92 | 95.92 | 95.92 | 95.92 | 95.92 | 95.92 | 95.92 | 95.92 | 96.33 | 96.33 |
| Bird 2 | 79.03 | 90.32 | 95.97 | 96.37 | 97.18 | 97.18 | 97.58 | 97.58 | 98.79 | 98.79 | 98.79 | 98.79 |
| Butterfly 2 | 35.19 | 86.36 | 95.96 | 97.14 | 97.14 | 99.16 | 99.49 | 99.66 | 99.66 | 99.66 | 99.66 | 99.66 |
| Fish 2 | 45.92 | 84.69 | 89.8 | 91.84 | 91.84 | 91.84 | 96.94 | 96.94 | 97.96 | 97.96 | 98.98 | 98.98 |

**Fig. 3.**
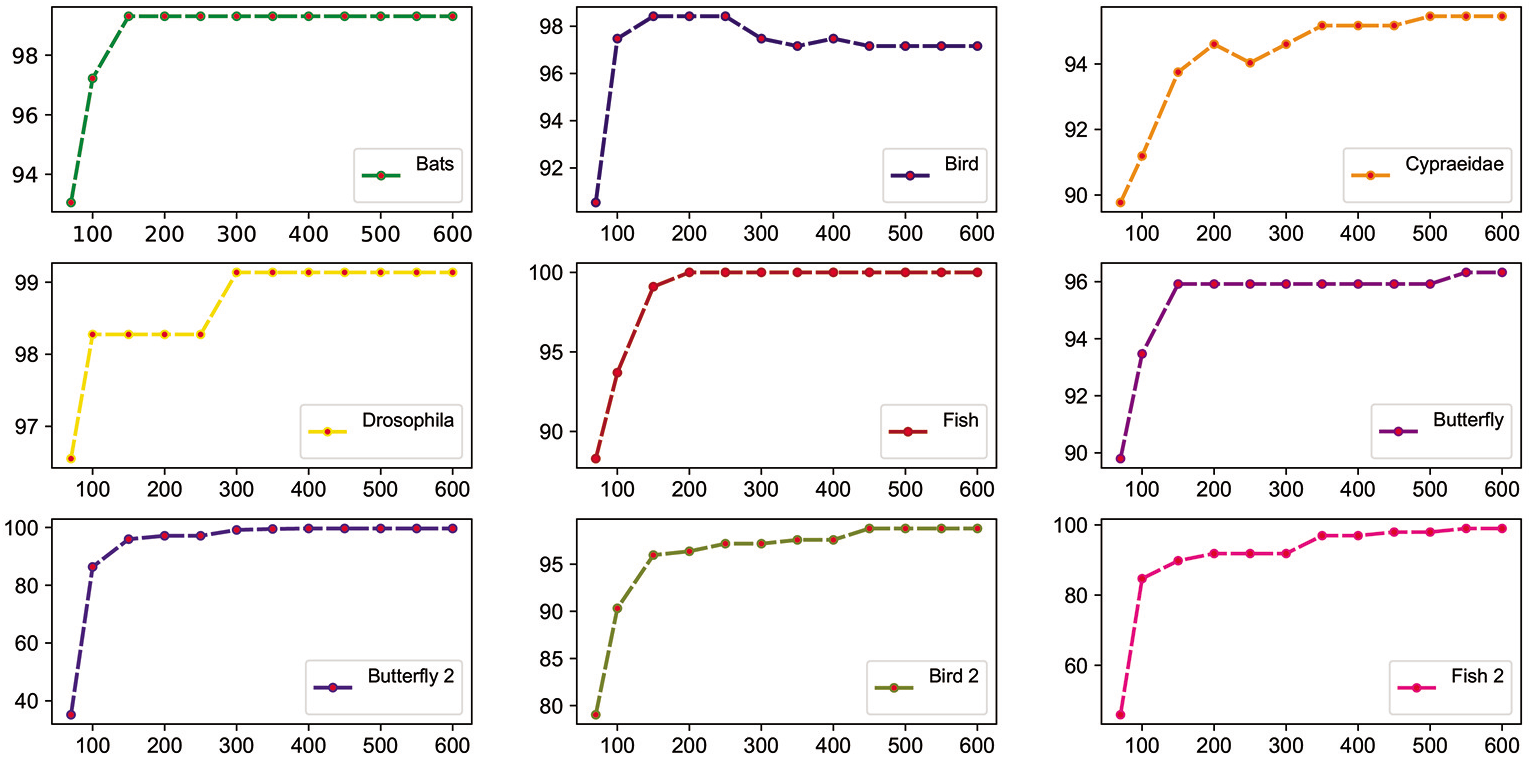
Lengthwise accuracy comparison for COI-based empirical datasets. From the lengthwise accuracy variations of each dataset, it is apparent that the accuracy is relatively consistent in the 200 bp-600 bp range but considerably falls below 200 bp.

**Fig. 4.**
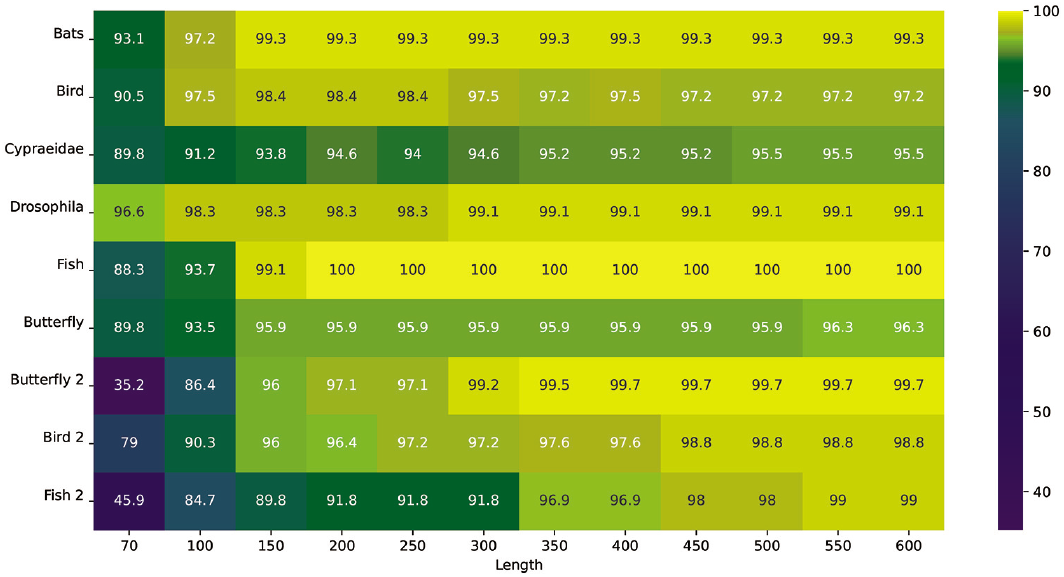
Heatmap of lengthwise accuracy comparison for COI-based empirical datasets. The lengthwise accuracy variation of Figure 3 is depicted in this heatmap. It is observed from the lengthwise color variation at each row that, 200 bp-600 bp provides a relatively consistent accuracy.

This similar approach was applied for the simulated datasets presented in Table III maintaing the same upper bound of 600 bp and lower bound of 70 bp. The lengthwise accuracy responses for the simulated datasets are presented in Table VII and demonstrated in Figure 5. From Figure 5 it is evident that, the accuracy score dropped insignificantly and steadily with the reduction of sequence length; however, it dropped significantly when the sequence length was reduced to < 200 bp.

**TABLE VII.** Accuracy Responses in the Simulated Datasets.

| Datasets | Lengths |  |  |  |  |  |  |  |  |  |  |  |
| --- | --- | --- | --- | --- | --- | --- | --- | --- | --- | --- | --- | --- |
|  | 70 | 100 | 150 | 200 | 250 | 300 | 350 | 400 | 450 | 500 | 550 | 600 |
| Ne1000 | 75.91 | 82.63 | 87.52 | 90.34 | 91.88 | 93.21 | 94.06 | 94.84 | 95.26 | 95.66 | 96.02 | 96.32 |
| Ne10000 | 81.05 | 86.85 | 91.28 | 93.21 | 94.43 | 95.14 | 95.54 | 95.84 | 96.08 | 96.26 | 96.46 | 96.60 |
| Ne50000 | 81.62 | 85.80 | 88.65 | 90.14 | 91.09 | 91.56 | 91.99 | 92.26 | 92.55 | 92.73 | 92.92 | 93.09 |

**Fig. 5.**
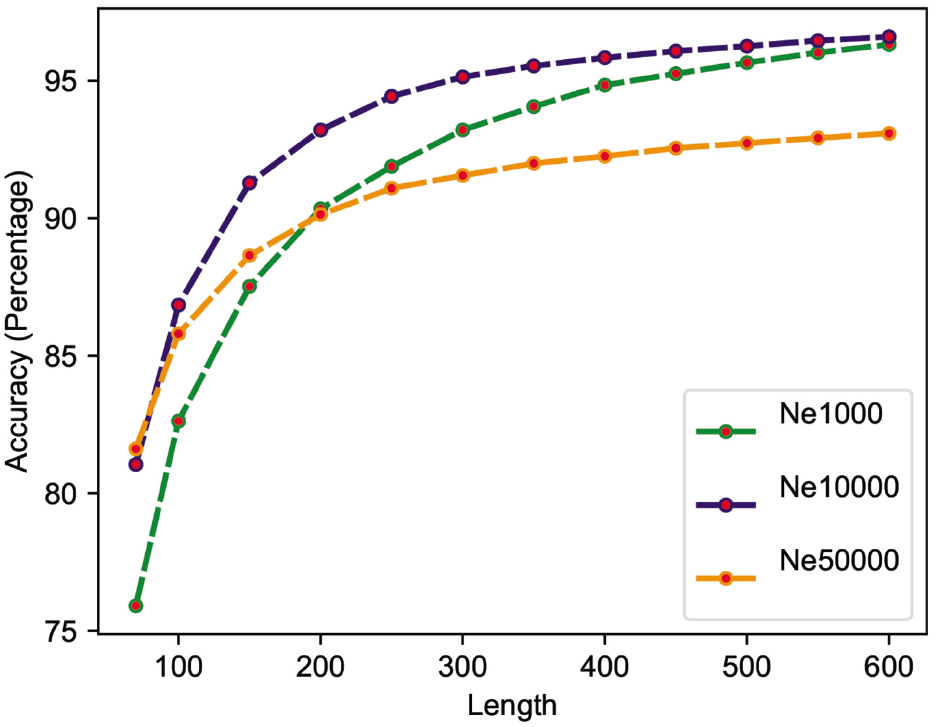
Lengthwise accuracy comparison for simulated datasets. This figure suggests a gradual but minor drop in the accuracy score when reducing the sequence length from 600 bp to 200 bp whereas a comparatively significant fall is noticed when the sequence length is reduced below 200 bp.

To obtain a complete overview of the effect of length on the species identification process, the datasets presented in Table III, based on the ITS and rbcL genes, were analyzed. For the datasets *Fungi* and *Algae*, we set the upper bound and lower bound for varying sequence lengths to 850 bp and 100 bp, respectively. For the dataset *Inga*, we set 1800 bp and 1050 bp as the respective upper and lower bounds because the full lengths of the barcode sequences in this dataset were > 1800 bp. In all three datasets, we decreased the length stepwise by 50 bases. The complete lengthwise average accuracy analysis of these datasets is presented in Table VIII. This analysis is also depicted in Figure 6.

**TABLE VIII.** Lengthwise Accuracy Responses in Fungi, Inga, and Algae Datasets.

|  |  | Lengths |  |  |  |  |  |  |  |  |  |  |  |  |  |  |  |
| --- | --- | --- | --- | --- | --- | --- | --- | --- | --- | --- | --- | --- | --- | --- | --- | --- | --- |
| Fungi | bp | 100 | 150 | 200 | 250 | 300 | 350 | 400 | 450 | 500 | 550 | 600 | 650 | 700 | 750 | 800 | 850 |
|  | % | 80.56 | 88.89 | 88.89 | 88.89 | 88.89 | 88.89 | 88.89 | 88.89 | 88.89 | 88.89 | 88.89 | 88.89 | 88.89 | 88.89 | 88.89 | 88.89 |
| Inga | bp | 1050 | 1100 | 1150 | 1200 | 1250 | 1300 | 1350 | 1400 | 1450 | 1500 | 1550 | 1600 | 1650 | 1700 | 1750 | 1800 |
|  | % | 78.79 | 78.79 | 78.79 | 78.91 | 85.09 | 86.63 | 87.53 | 87.40 | 87.40 | 87.40 | 87.14 | 89.71 | 90.48 | 90.10 | 95.11 | 95.11 |
| Algae | bp | 100 | 150 | 200 | 250 | 300 | 350 | 400 | 450 | 500 | 550 | 600 | 650 | 700 | 750 | 800 | 850 |
|  | % | 90.0 | 90.0 | 90.0 | 90.0 | 100.0 | 100.0 | 100.0 | 100.0 | 100.0 | 100.0 | 100.0 | 100.0 | 100.0 | 100.0 | 100.0 | 100.0 |

**Table Notes:** In this table, accuracy responses are demonstrated at different lengths for each of the Fungi, Inga and Algae datasets where respective results for each dataset consist of two rows. For each dataset, the first row indicates the lengths of the sequence (denoted by bp) and the second row indicates the corresponding accuracies (denoted by %).

**Fig. 6.**
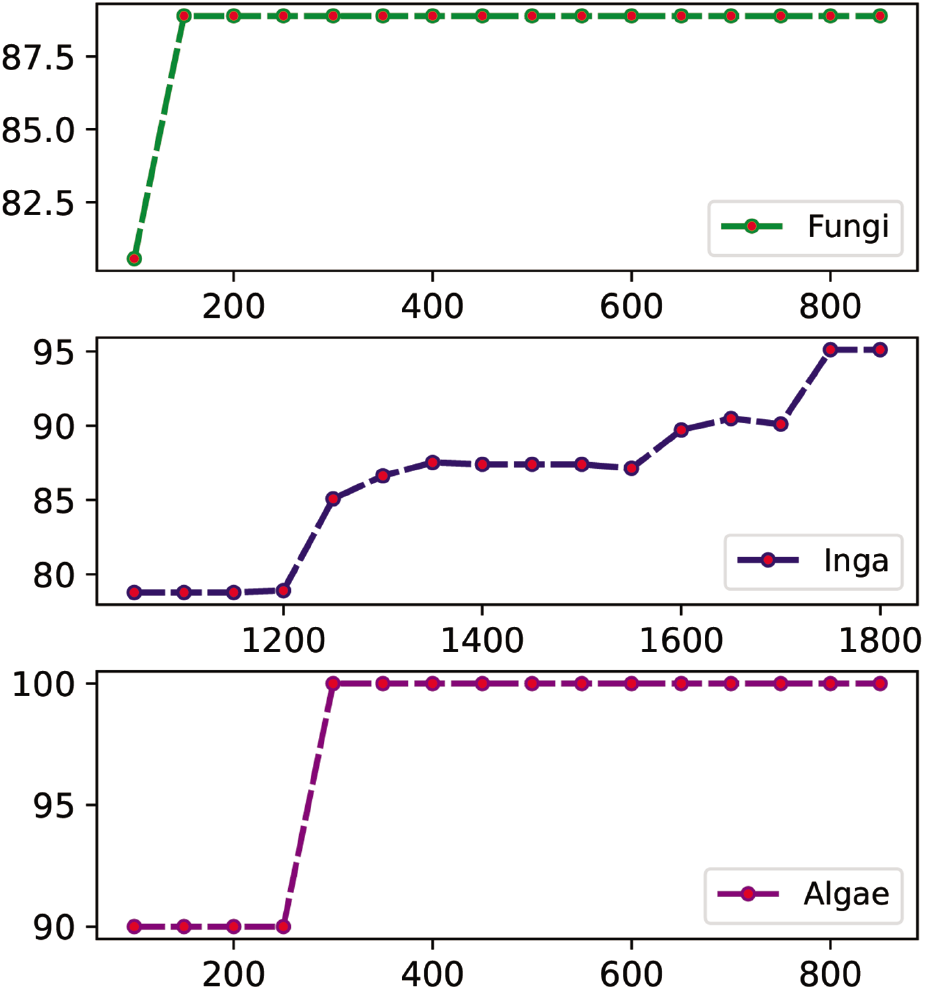
Lengthwise accuracy comparison in Fungi, Inga and Algae datasets. This figure demonstrates a sharp fall in accuracy scores for Algae and Fungi dataset around sequence lengths below 200 bp-250 bp. The length and subsequent length wise accuracy responses in Inga dataset are not similar to those of Fungi and Algae datasets, but still a decline in accuracy score can be observed with the gradual decrease of sequence length.

Figure 6 shows that although the contexts differed from the previous lengthwise accuracy patterns of COI-based empirical datasets and simulated datasets, *Algae* and *Fungi* showed a sharp fall in accuracy scores for sequence lengths below 200-250 bp. However, this corroboration cannot be extended to *Inga* because its length-bounds were selected differently. However, it still showed a decrease in accuracy similar to the others when the lengths were gradually decreased from full length. To compare these three datasets on a common ground, average accuracy responses were calculated at various percentage levels of their full lengths. The results are presented in Table IX and depicted in Figure 7. No persistent range similar to the previous COI-based empirical datasets were evident from these representations of ITS and rbCL-based datasets. However, the lengthwise accuracy of *Fungi* and *Algae* decreased sharply at approximately 20 %-30 % of their full lengths whereas the accuracy in the *Inga* dataset considerably decreased initially at around 70 % and later at 40 % of its full length. For COI-based and simulated datasets, the length (~200 bp) below which a sharp decline was observed was approximately 30 % of the full length of barcode sequence. In Table VIII and Table IX, for each dataset the first row presents the sequence lengths and the second row presents the corresponding accuracy scores.

**TABLE IX.** Accuracy Responses in Fungi, Inga, and Algae Datasets at Different Length Percentages

|  |  | Percentages of Lengths |  |  |  |  |  |  |  |  |  |
| --- | --- | --- | --- | --- | --- | --- | --- | --- | --- | --- | --- |
|  |  | 10% | 20% | 30% | 40% | 50% | 60% | 70% | 80% | 90% | 100% |
| Fungi | bp | 93 | 186 | 279 | 372 | 465 | 558 | 651 | 744 | 837 | 930 |
|  | % | 79.44 | 93.33 | 93.33 | 93.33 | 93.33 | 93.33 | 93.33 | 93.33 | 93.33 | 93.33 |
| Inga | bp | 183 | 367 | 551 | 735 | 919 | 1102 | 1286 | 1470 | 1654 | 1838 |
|  | % | 46.33 | 66.84 | 67.09 | 76.41 | 76.54 | 79.09 | 87.57 | 88.49 | 90.06 | 94.96 |
| Algae | bp | 112 | 225 | 338 | 451 | 564 | 676 | 789 | 902 | 1015 | 1128 |
|  | % | 90.0 | 90.0 | 100.0 | 100.0 | 100.0 | 100.0 | 100.0 | 100.0 | 100.0 | 100.0 |

**Table Notes:** In this table, accuracy responses are demonstrated at different percentage levels of the full sequence lengths for each of the Fungi, Inga and Algae datasets where respective results for each dataset consist of two rows. For each dataset, the first row indicates the lengths of the sequence (denoted by bp) at the mentioned percentage levels and the second row indicates the corresponding accuracies (denoted by %).

**Fig. 7.**
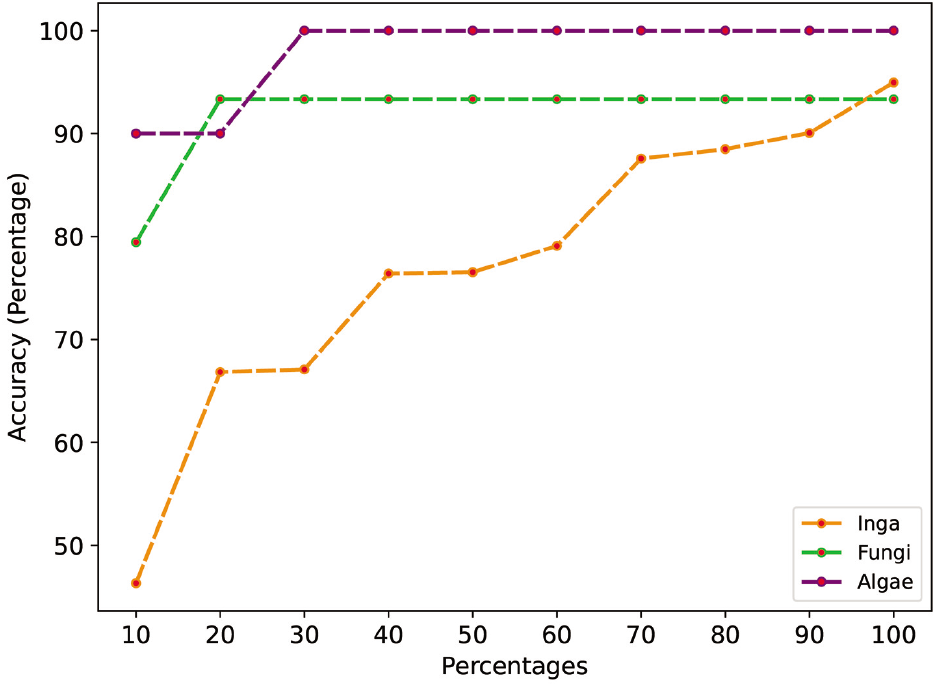
Accuracy responses in Fungi, Inga and Algae datasets at different length-percentages. As the sequence lengths of these three datasets are different, average accuracy responses were further calculated at various percentage levels of their full lengths to bring uniformity.

To support our findings, Figure 8 was adapted and redrawn from the work of [15]. In this figure, the proportion of species resolution with different DNA barcode size is depicted. The shapes in the individual plots of Figure 3 match the shape in Figure 8 implying the consistency between these separate but similar findings. From other related studies regarding mini-barcodes, it was observed that full length barcodes provide marginally better identification success rates than mini-barcodes with 200-400 bp sequence length, whereas the identification ability is significantly compromised only if the barcode length falls below 200 bp [3]. From the present study findings, an approximation of 200-350 bp mini-barcode length is proposed for use in species identification using the DNA barcode approach with a supervised machine learning model because it has a significant possibility of performing with a reasonably accurate score.

**Fig. 8.**
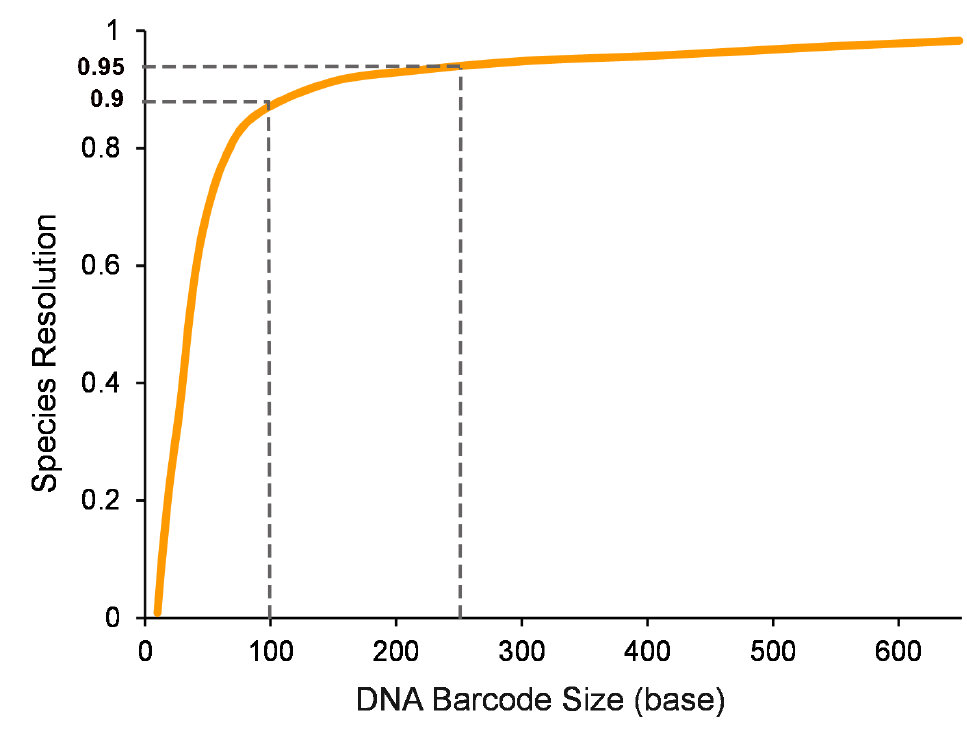
Change of species resolution in different DNA mini-barcode lengths (adapted and redrawn from [15]). In the demonstration of [15], it is asserted that although best performance can be achieved at full length barcode with 97 % species resolution, 100 bp and 250 bp mini barcode provide 90 % and 95 % identification success respectively.

## VI. Conclusion

In the present study, we provide an approximation of the suitable length of DNA mini-barcodes from a supervised machine learning approach by analyzing the lengthwise accuracy during the species identification process. A Naïve Bayes classifier was used to determine the effect of varying DNA barcode lengths on identification accuracy. Our findings from this machine learning perspective are consistent with those of related biological works where DNA mini-barcodes were used and analyzed to aid the species identification process. Although we proposed an approximate effective length of minibarcodes for *COI* gene based DNA sequences, further studies are required for plants and *ITS* gene barcode based organisms. Improving the quality of the sequence alignment using suitable multiple sequence alignment tools/techniques would improve the identification accuracy. Other machine learning algorithms should also be applied to enrich the comprehensiveness of the findings.

## Acknowledgment

We sincerely thank Dr. Md. Shamsuzzoha Bayzid for his valuable guidelines and immense support during this study.

## Footnotes

1 *DNA Barcodes classification with supervised machine learning techniques*, Institute for Systems Analysis and Computer Science, Italian National Research Council. Accessed: Aug. 2021. [Online]. Available: dmb.iasi.cnr.it/ supbarcodes.php

2 *Research Challenges Related to Data Analysis and the DNA Barcode of Life Initiative: Data Sets*, Center for Discrete Mathematics and Theoretical Computer Science, May 2007. Accessed: Aug. 2021. [Online]. Available: http://dimacs.rutgers.edu/archive/Workshops/BarcodeDataSets/

3 Public data set “DS-BUNEACAR”, BOLD Systems. Accessed: Aug. 2021. [Online]. doi: dx.doi.org/10.5883/DS-BUNEACAR.

4 M. Karim, R. Abid. “DNA_Barcode.” Github.com. Accessed: Aug. 2021. [Online]. Available: https://github.com/MohimenulRafi/DNA_Barcode

